# ORFik: a comprehensive R toolkit for the analysis of translation

**DOI:** 10.1101/2021.01.16.426936

**Authors:** Håkon Tjeldnes, Kornel Labun, Yamila Torres Cleuren, Katarzyna Chyżyńska, Michał Świrski, Eivind Valen

## Abstract

**Background:** With the rapid growth in the use of high-throughput methods for characterizing translation and the continued expansion of multi-omics, there is a need for back-end functions and streamlined tools for processing, analyzing, and characterizing data produced by these assays.

**Results:** Here, we introduce ORFik, a user-friendly R/Bioconductor toolbox for studying translation and its regulation. It extends GenomicRanges from the genome to the transcriptome and implements a framework that integrates data from several sources. ORFik streamlines the steps to process, analyze, and visualize the different steps of translation with a particular focus on initiation and elongation. It accepts high-throughput sequencing data from ribosome profiling to quantify ribosome elongation or RCP-seq/TCP-seq to also quantify ribosome scanning. In addition, ORFik can use CAGE data to accurately determine 5’UTRs and RNA-seq for determining translation relative to RNA abundance. ORFik supports and calculates over 30 different translation-related features and metrics from the literature and can annotate translated regions such as proteins or upstream open reading frames. As a use-case, we demonstrate using ORFik to rapidly annotate the dynamics of 5’ UTRs across different tissues, detect their uORFs, and characterize their scanning and translation in the downstream protein-coding regions.

**Availability:** http://bioconductor.org/packages/ORFik

## Background

Messenger RNAs (mRNAs) can be divided into three regions: the transcript leader sequence also known as the 5’ untranslated region (5’ UTR), the coding sequence (CDS), and the trailer sequence or 3’ untranslated region (3’ UTR). Translation is normally initiated by the binding of the 43S ribosome to the mRNA, adjacent to the cap. The 43S preinitiation complex, consisting of the small ribosomal subunit (SSU) and eukaryotic initiation factors (eIFs), then proceeds to scan downstream, until it reaches a favorable initiation context at the start of an open reading frame (ORF). Here, it recruits the 60S large ribosomal unit, which together with the 43S forms the 80S elongating complex. The 80S, then, proceeds to translate the ORF, processing it codon-by-codon, until it reaches a terminating codon and the protein synthesis is complete^1^.

While eukaryotic transcripts typically encode only a single protein, evidence from high-throughput methods has revealed that many 5’ UTRs contain short upstream ORFs (uORF) that can be translated^2^. While the functional importance of uORFs is still debated, several uORFs have been found to regulate gene expression^2,3^. This primarily occurs by hindering ribosomes from reaching the protein-coding ORF leading to translational inhibition. This demonstrates that at least a subset of uORFs is functionally important.

While translation was previously studied on a gene-by-gene basis, the introduction of ribosome profiling (ribo-seq) and later, translation complex profiling (TCP-seq) and ribosome complex profiling (RCP-seq) has made it possible to obtain a snapshot of translating and scanning ribosomes across the whole transcriptome^4,5,6^. Together with information on RNA levels and isoforms, protein-coding ORFs and uORFs can be identified and their translational levels can be quantified. Getting functional insight from sequencing data requires robust computational analysis. Ribo-seq, being a mature assay, has a number of software packages and web services designed specifically to handle it. TCP-seq^5,7,8^ and RCP-seq^6^, on the other hand, are much less supported. These methods need tools that can consider both 43S and 80S dynamics as well as the relationship between these.

Another complicating factor is that many genes have alternative transcription start sites (TSSs)^9^. For the study of translation initiation, accurate annotation of 5’ UTRs is required, otherwise, it will be challenging to determine which uORF candidates should be included in the analysis. The choice of TSS dictates which uORFs are present in the 5’ UTR. In certain cases, uORFs are only present in specific tissues with the correct variant of the 5’ UTR^10^.

To address these challenges and provide a comprehensive tool for studying translation in custom regions we developed ORFik, a Bioconductor software package that streamlines the analysis of translation. It supports accurate 5’ UTR annotation through RNA-seq and cap analysis of gene expression (CAGE), detection and classification of translated uORFs, characterization of sequence features, and the calculation of over 30 features and metrics used in the analysis of translation (Table S3).

## Implementation

ORFik is implemented as an open-source software package in the R programming language, with C++ implementations for the possible bottlenecks in large datasets. While tools for certain steps of translation analysis exist, none of them support analysis of TCP-seq / RCP-seq in combination with CAGE and many of these are either; online tools^11^, or are limited to studying only specific steps or aspects of translation^12^. A full comparison between related tools can be found in Table S6^13,13,14,15,16,17–20,21,22,23^.

ORFik is highly optimized and fast. To achieve this, we have reimplemented several functions in the Bioconductor core package GenomicFeatures that are slow for larger datasets, like converting from transcript coordinates to genomic coordinates and vice versa. In addition, to aid with the ever-increasing size of datasets, we have focused on allowing faster computation of large bam files with our format “.ofst” based on the Facebook compression algorithm zstd^24,25^. “.ofst” is a serialized format (see the section on Optimized File Format), with optional collapsing of duplicated reads, enabling near-instantaneous data loading (Table S1).

## Overview

A typical workflow takes transcriptome/genome annotation and high-throughput sequencing data as input, processes these to make transcriptome-wide tracks (Figure 1A), and use these to either make summary statistics for all genes or transcripts or to characterize one or more specific transcriptomic regions. Regions can be of any type and are completely user-customizable, but typically consist of genes, 5’ UTRs, CDSs, uORFs, start codons, or similar. ORFik can then be used to calculate summary statistics and features for all candidate regions.

**Figure 1:**
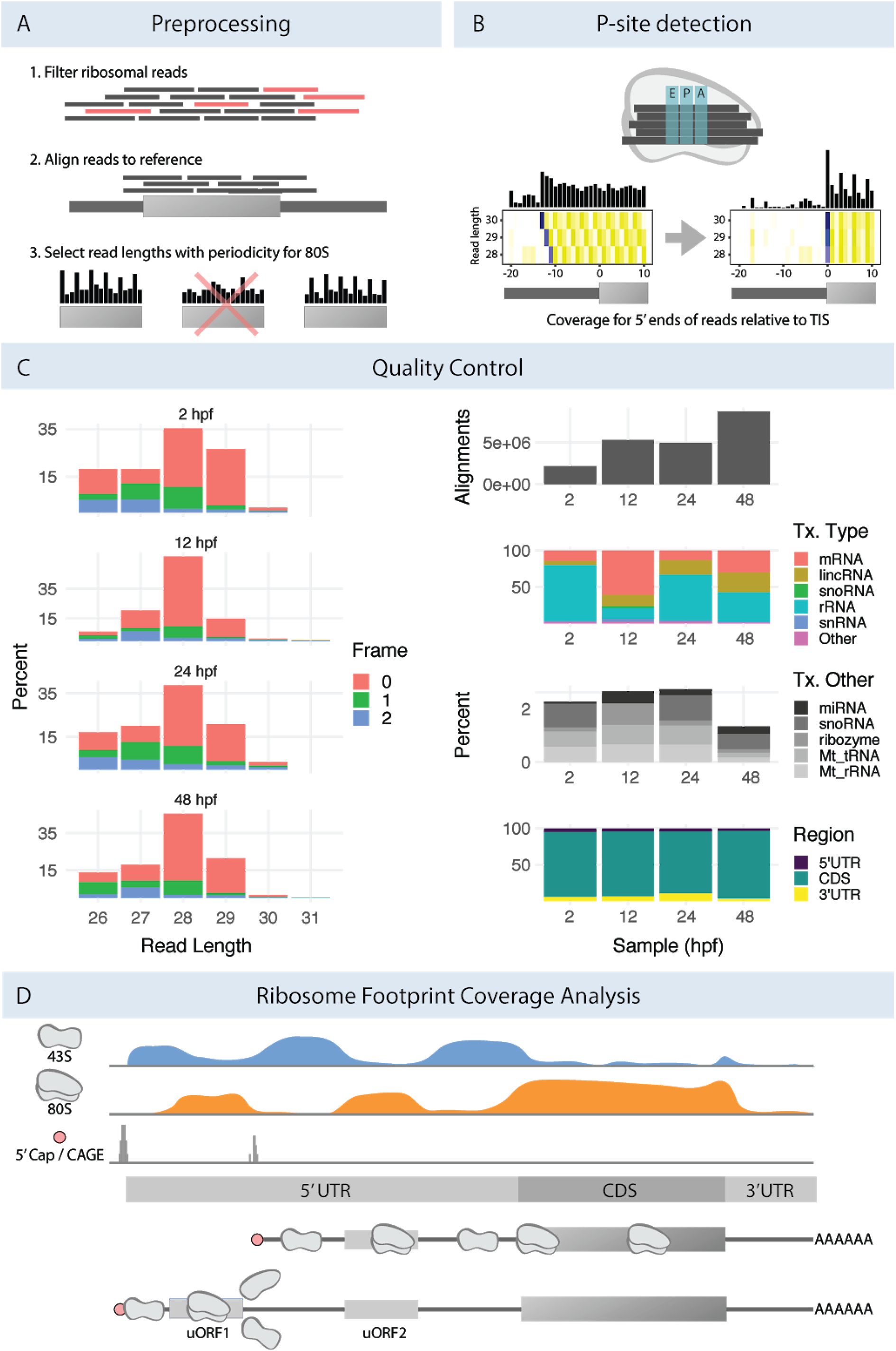
ORFik functionality. A) ORFik supports a number of preprocessing steps including 1) removal of rRNA contamination) and other ncRNA), 2) Alignment to genome or transcriptome, 3) selecting 80S read lengths that display periodicity consistent with translation. B) To identify the P-sites for reads (top illustration) from an 80S library ORFik performs change point detection on the 5’ ends of reads across a window over translation tnitiation sites. This determines the location of reads from initiating ribosomes and the distance from their 5’ end to the start codon. This is done separately for each read length (heatmap). C) Examples of figures used to perform quality control on the data from four stages of zebrafish development^26^. Left column: The percentage of reads in each translation frame over CDSs after P-shifting, stratified by read-length. Right column: 1) Number of aligned reads after filtering contaminants. 2) Percentage of reads aligning to various transcript types (>1%). 3) Percentage alignment of each transcript type in the “Other” group. 4) Percentage of reads aligning to mRNA falling into the CDS or UTRs. For more QC examples see Figure S4 and S5. D) Read coverage tracks for 43S ribosomal small subunit, 80S translating ribosomes and CAGE-defined transcription start sites (TSS) for a model gene. The gene has two transcript isoforms with different TSSs and 5’UTRs. The second isoform harbors an additional uORF leading to translational repression through ribosome dissociation (uORF 1). ORFik can assist in detecting such differences in isoform through differential expression, visualization, and metrics.

ORFik supports standard translation analysis: it can map reads from RNA-seq and ribo-seq, it performs trimming and P-site shifting of ribo-seq reads quantifies ribosomal occupancy (Figure 1B), characterizes ORFs, and creates a range of plots and predictions. In addition, it supports the analysis of translation initiation through TCP-seq, RCP-seq, and CAGE. It can quantify translation initiation through scanning efficiency and ribosome recruitment and can correlate these with sequence elements. Overall, ORFik provides a toolbox of functions that is extremely versatile and enables the user to go far beyond standardized pipelines.

## Obtaining and preprocessing data

The first step in ORFik is obtaining and preprocessing data (Figure 1A). It automates direct download of datasets from the NGS repositories: SRA^27^, ENA^28^, and DRA^29^, download of annotations (FASTA genome and GTF annotation) through a wrapper to biomartr^30^, while also supporting local NGS datasets and annotations. The sequencing data can be automatically trimmed with fastp^31^ with auto removal of all possible adapters or using presets for the most common Illumina adapters. Following this, ORFik can detect and remove any of the known contaminants (e.g. Phi X 174, rRNA, tRNA, non-coding RNAs) and finally align the reads using STAR^32^.

### Quality control and track creation

Post-alignment quality control is initiated by calling the function ORFikQC. ORFik will output quality control plots and tables for comparisons between all runs in an experiment and the whole process is streamlined to be as fast and simple as possible for the user (Figure 1C). Among others, plots of the meta coverage across all transcripts regions and correlation plots between all pairs of samples (Figure 1D, S4) are generated. Furthermore, essential mapping statistics are calculated such as the fraction of reads mapping to important features, contaminants, and transcript regions (Figure 1C).

### Single-base resolution of transcription start sites

Many organisms have extensive variations in the use of isoforms between tissues and can often use a range of sites to initiate transcription. Since many analyses of translation depend on an accurate annotation of 5’UTRs, ORFik enables the use of CAGE data in its pipeline. CAGE is a high-throughput assay for the precise determination of transcription start sites (TSSs) at single-base pair resolution^33^. ORFik makes use of CAGE (or similar 5’ detection assays) to reannotate transcripts. In the standard pipeline (Figure S1), ORFik identifies all CAGE peaks in promoter-proximal regions and assigns the largest CAGE peak as the TSS. This can be customized to only consider specific thresholds or exclude ambiguous TSSs that are close to the boundary of other genes. An example of analysis performed on CAGE-reassigned TSSs can be seen in Figure 3B.

**Figure 2:**
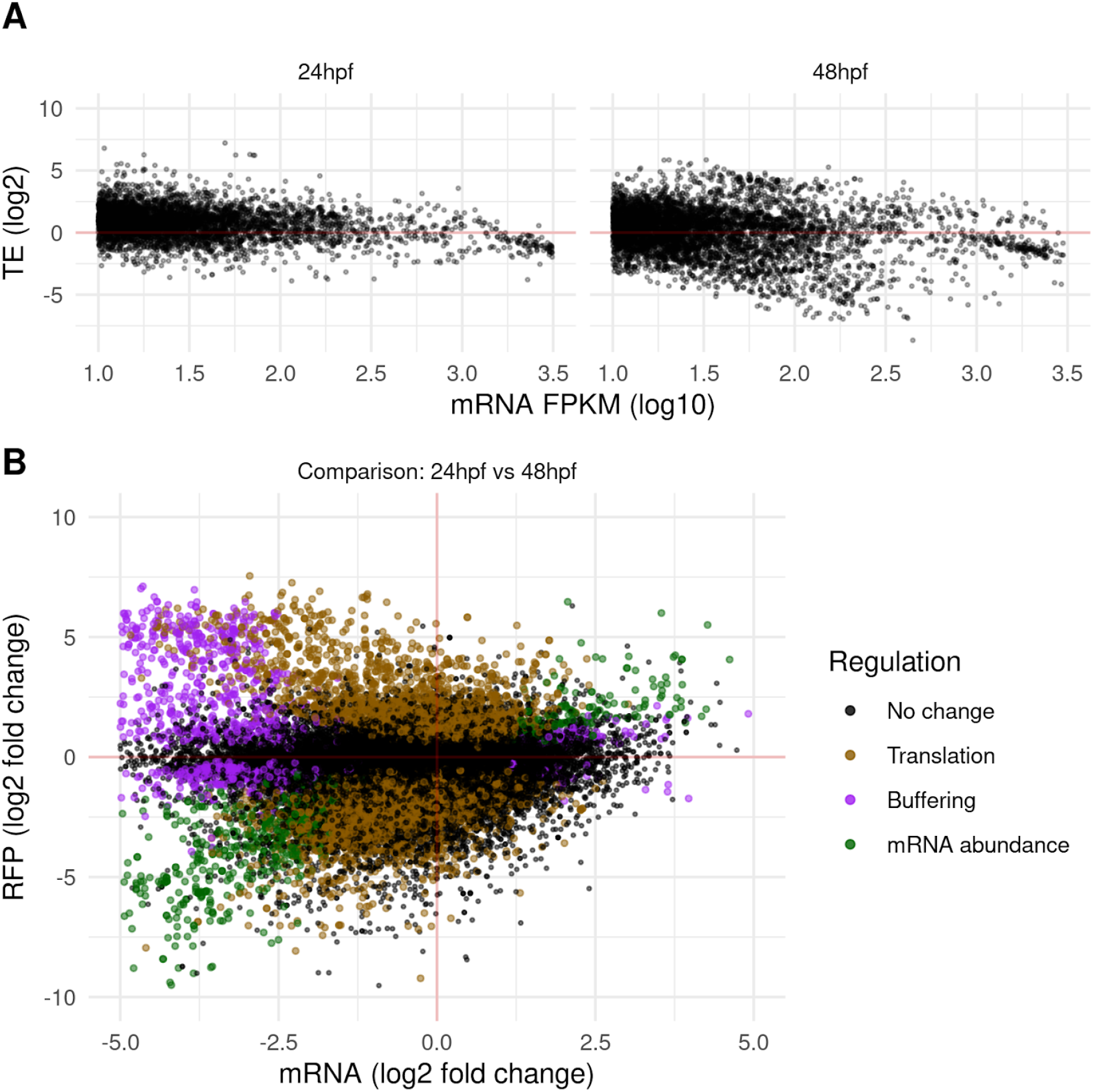
Analysis of translation. ORFik supports within-group (A) and between-group (B) translation analysis plots. Example data from two zebrafish developmental stages (described later): 24 hpf and 48 hpf, each with 2 replicates^26^. A) TE vs mRNA levels using the average values across replicates. B) Differential translation analysis between conditions. Non-significant genes are colored black while significant genes (p<0.05) are grouped into 3 categories according to deltaTE classification^41^: Translational regulation — mRNA abundance is static while translation changes (orange), Buffering — mRNA, and ribosome profiling regulated in opposite directions (purple), mRNA abundance regulation — change in mRNA abundance with a corresponding change in ribo-seq levels (green).

**Figure 3:**
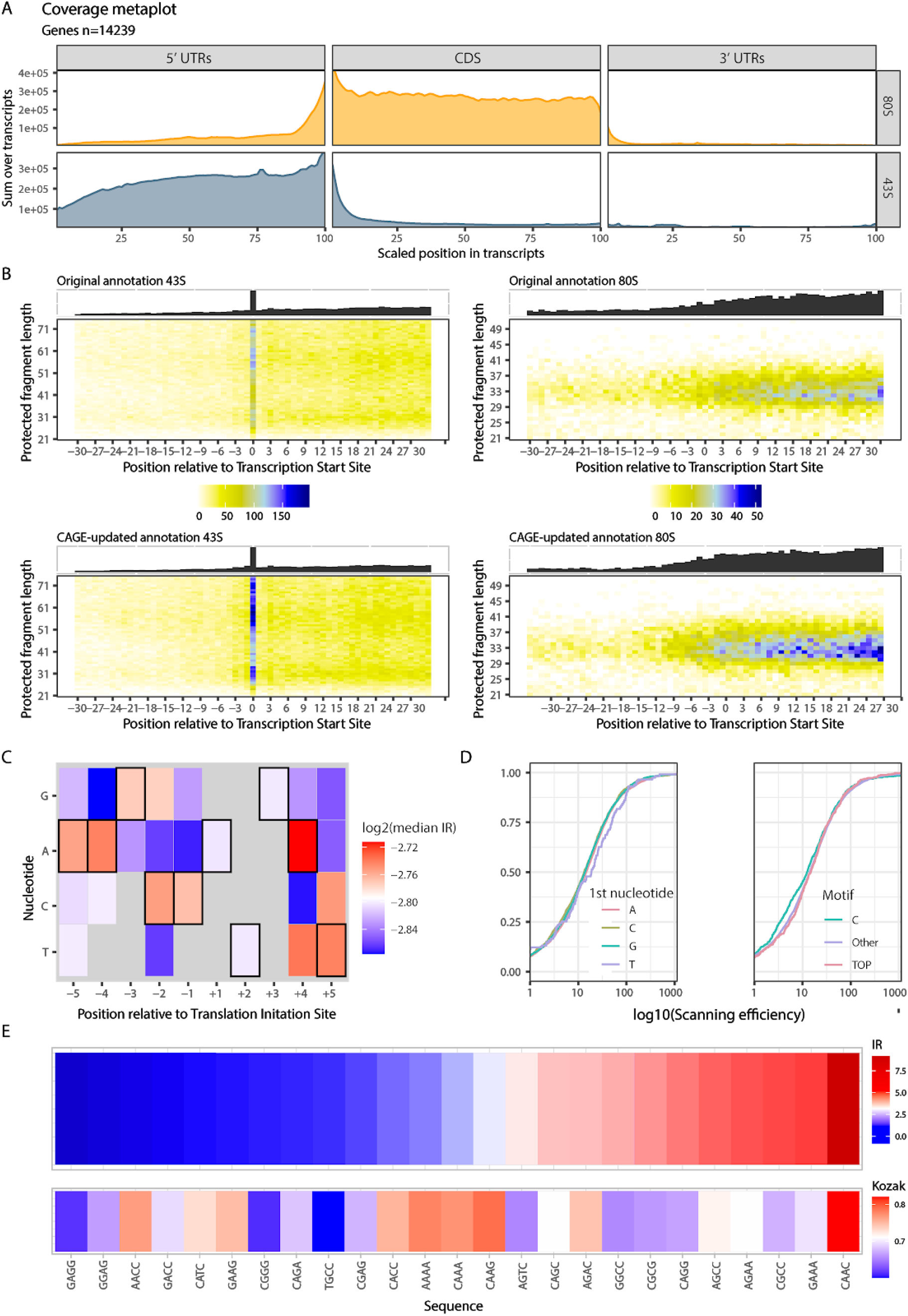
Analysis of translation initiation. A) Meta coverage plot of RCP-seq data over 5’ UTRs, CDSs, and 3’ UTRs from all mRNA transcripts^8^. Regions are scaled io be the same size and coverage is displayed as the sum of reads for the translating (80S) and scanning ribosomes (43S). B) 5’ end coverage heatmaps of 43S reads (left column) and 80S (right column) from TCP-seq relative to transcription start site. Coverage is shown before (upper row) and after (lower row) reannotation of transcription start sites with CAGE data. C) Analysis of initiation context showing the median initiation rate (IR) for translation initiation sites with a specific base (x-axis) at a specific position (y-axis). The strongest consensus is displayed with a black frame. D) Analysis of ribosome recruitment showing the relationship between different TSS contexts and 43S scanning efficiency. Left: empirical cumulative distribution function (ecdf) of scanning efficiency colored by the first nucleotide in the 5’UTR. Right: ecdf of scanning efficiency tor 3 different motifs; a C nucleotide, the TOP motif (C, then 4 pyrimidines), and all other TSS variants. E) initiation contexts ranked by IR. Top: mean IR for all CDSs with specified sequence (x-axis) relative to TIS (−4 to −1). Bottom: similarity score to human Kozak sequence as defined in Grzegorski et al. 2014^47^.

### Automatic read length determination

When performing ribo-seq it is customary to size-select a range of fragments that correspond to the size protected by a single ribosome. This is because not all isolated fragments necessarily originate from regions protected by ribosomes. Instead, these can be the product of other sources such as RNA structure, RNA binding proteins or incomplete digestion^34^ This filtering of sizes is typically performed first in the lab and then computationally. To identify which read lengths most likely originate from actual ribosome protected footprints (RFPs), ORFik identifies read lengths that display 3 nucleotides (nt) periodicity over protein-coding regions, indicative of ribosome translocation (Figure 1A). For each fragment length, we sum the 5’ ends of footprints mapping to the first 150 nucleotides in the CDSs of the top 10% of protein-coding genes ranked by coverage and keep lengths with at least 1000 reads. The resulting vectors are subject to discrete Fourier transform, and the fragment lengths whose highest amplitude corresponds to a period of 3 are considered to be bonafide RFPs (Figure S6). By default, all read lengths that lack this periodicity are filtered out^35^.

### Sub-codon resolution through P-shifting

Sequenced RFP reads span the whole region where the ribosome was situated. In many analyses, however, it is interesting to increase the resolution and determine exactly which codon the ribosome was processing when it was captured, the so-called P-site. Several methods of determining P-site location within footprints (P-shifting) have previously been developed^35, 36, 12, 23^. ORFik predicts the P-site offset per read length, taking inspiration from the Shoelaces algorithm^35^. To determine the P-site offset in the protected footprints, ORFik considers the distribution of the different read lengths over the TIS region. For each read length ORFik takes the 5’ end of all reads from all genes and sums these for each coordinate relative to the TIS. ORFik then performs a change point analysis to maximize the difference between an upstream and downstream window relative to the changepoint (Figure S7). This analysis results in the most probable offset to shift the 5’ ends of fragments protected by initiating ribosomes exactly at the TIS (Figure 1B). The function can process any number of libraries and can supply log files and heatmaps over the start and stop codon used to verify that the P-site detection was correct (Figure S2) and users can manually override the suggested shifts. The user can choose between several formats, where the 2 default formats are wiggle format (wig) and ofst (for very fast loading into R).

## Analysis

After preprocessing the user creates an instance of the “experiment” class. This summarizes and provides information about the data and makes it possible to analyze, plot, or create features for any number of NGS libraries. Experiments contain all relevant metadata so that naming and grouping in plots can be performed automatically. It also specifies the correct annotation and genomic sequence files for the data, enabling automatic downloading of these. To facilitate sharing between researchers the experiment class is constructed to not contain any local information like file paths. This makes it easy to send a short single script to collaborators that can perfectly replicate the entire process starting from scratch and producing identical downstream analyses and plots.

### ORFik extends GenomicRanges and GenomicFeatures to the transcriptome

To facilitate the analysis of any region in the transcriptome ORFik extends the functionality of two Bioconductor core packages: GenomicRanges and GenomicFeatures^24^. The intention of the first package was to provide robust representation and facilitate manipulation of contiguous (non-spliced) genomic intervals. Genomic Features, on the other hand, aimed at extending this functionality to spliced ranges and more complicated annotations, like gene models. The high-speed functions for intra- and inter-ranges manipulation of Genomic Ranges are, however, inconsistent or completely lacking for the spliced ranges. ORFik extends these packages with more than 50 new functions for spliced ranges and more effectively converts between genomic and transcriptomic coordinates (Table S1). ORFik also supports fully customizable subsets of transcript regions and direct subsetting of among others start codons, stop codons, transcription start sites (TSS), translation initiation sites (TIS) and translation termination sites (TTS).

ORFik also contains functions to facilitate rapid calculation of read coverage over regions. Beyond basic coverage per nucleotide, ORFik supports different coverage summaries like read length or translation frame in ORFs (Table S2). All coverage functions also support the input reads as collapsed reads (merged duplicated reads with a meta column describing the number of duplicates). This greatly speeds up calculations and reduces memory consumption, especially for short-read sequencing data characterized by high duplication level (e.g. ribo-seq).

### Visualizing meta coverage

Metaplots are a useful way of visualizing read coverage over the same relative region across multiple transcripts. ORFik implements these plots through a generalized syntax used for intuitive one-line functions. All plots can be extended or edited as ggplot objects (R internal graphic objects^37^). Since meta coverage and heat maps can be represented in multiple ways that emphasize different features of the data ORFik provides 15 different data transformations for metaplots (e.g. raw sum, mean, median, transcript normalized, position normalized, mRNA-seq normalized) (Table S2). It also provides filters to avoid bias from single occurrences such as extreme peaks caused by contaminants in the data (e.g. non-coding RNAs). ORFik can filter these peaks or whole transcripts, to make plots more representative of the majority of the regions.

### Identifying and characterizing open reading frames

ORFs can be split into a hierarchy based on the available evidence: 1) sequence composition and presence in a transcript, 2) active translation, measured with ribo-seq, 3) peptide product detection 4) confirmation of function. ORFik addresses the two first levels of this hierarchy.

Obtaining all possible ORFs based on sequence is accomplished through a scan of user-provided FASTA sequences to find candidate ORFs. The search is efficiently implemented using the Knuth-Morris-Pratt^38^ algorithm with binary search for in-frame start codons per stop codon (Table S1). It supports circular prokaryotic genomes and fast direct mapping to genomic coordinates from transcript coordinates. The provided sequence data can be any user-provided files or biostrings objects but the most straightforward approach is to obtain these from ORFiks own data loading functions. After identification, ORFs can be saved (e.g. in BED format with color codes) and loaded into a genome browser like IGV or UCSC genome browser for visualization.

If based purely on the sequence, the number of ORFs in most genomes is vast^39^. When searching for uORFs or novel genes it can therefore be advantageous to move to the second step of the ORF hierarchy and also consider the ribosomal occupancy of the novel region. However, when analyzing small regions, like putative uORFs, simply observing the presence of reads will often have low predictive power. This stems from two issues: The first is a sampling problem, in that ribosomes over a short region might be transient and simply not get sampled and sequenced unless the occupancy is particularly high. The second is the aforementioned problem that not all reads originate from ribosomes. A weak ribo-seq signal alone is therefore not conclusive as evidence for translation^40^.

To address this, several metrics or features have been produced that quantify how RNA fragments that originate from ribosome protection behave over verified protein-coding regions. ORFik currently supports more than 30 of these metrics that have been previously described in the literature, by us and others (Table S3). ORFik also provides a wrapper function computeFeatures that produces a complete matrix of all supported metrics with one row per ORF or region. This resulting output matrix can be used to characterize specific ORFs or as input to machine learning classifiers that can be used to predict novel functional ORFs. By using a set of verified ORFs (e.g. known protein-coding sequences) the user can construct a positive set and use the features learned from them to classify the set of new candidate ORFs. Alternatively, the matrix may be exported and provided as input to other tools.

#### Differential expression

ORFik supports various ways of estimating differential expression or preparing data for other tools. For visualization, it supports standard plots for studying fold changes of translation and RNA and the relationship between expression and translational efficiency (Figure 2A). Since many tools require raw counts as input for analysis of expression^41,42,43^ ORFik can provide count data tables through the countTable function. This can optionally collapse and merge replicates or normalize data to FPKM values. These count tables can be supplied to DESeq2, anota2seq, and other tools to perform differential expression analysis^41,44^. Alternatively, an implementation of the deltaTE algorithm^45^, is also included in ORFik, which can detect translational regulation between conditions using a Wald test for statistical significance (Figure 2B).

## Results

To illustrate the functionality of ORFik we show two use cases, where we 1) study translation initiation with profiling of small ribosomal subunits and 2) make a simple pipeline to predict and characterize translated uORFs.

### Using ORFik to analyze translation initiation

When analyzing data that includes small subunit profiling the goal is often to understand or characterize the regulation of translation initiation. ORFik was here applied to a TCP-seq data set from HeLa cells^8^, and easily produces metaprofiles over transcripts, separated into 43S scanning and 80S elongating ribosomes (Figure 3A). As expected, the 43S scanning ribosomes display the highest coverage in 5’UTRs, while elongating ribosomes are enriched over the CDSs. To obtain accurate measurements of scanning over 5’UTRs, CAGE data was used to reannotate the TSSs. The effect on the coverage of 40S complexes around the TSS as a result of this reannotation can be viewed in Figure 3B. These heatmaps have been *transcript-normalized*, meaning that all the reads mapping to one transcript has been normalized to sum to 1. This has the effect of weighing each transcript equally, but ORFik supports a range of other normalization methods that emphasize different aspects of the data (Figure S3). The increased coverage and sharper delineation of the TSS that can be seen in these plots illustrate that CAGE-reannotation is important even in well-annotated transcriptomes like the human. Accurate TSS is also important when studying features at the start of the transcript such as the presence of a TOP motif^46^. TOP motifs are known to be involved in the regulation of specific genes and ORFik can correlate such features to other observations of the transcript. An example of such an analysis is associating motifs with the scanning efficiency (SE) — the number of scanning ribosomes on the 5’UTR relative to the RNA abundance^5^. In our latest study, we showed an association between ribosome recruitment and TOP motifs during early development^6^. ORFik easily produces a similar analysis for the HeLa cells revealing no such relationship in these cells.

Beyond recruitment, an important part of translation initiation is the recognition of the start codon. This is facilitated by the context surrounding it and ORFik provides several ways of exploring these features. At the level of basic visualization heatmaps, similar to those for TSS, can be produced to explore the conformations and periodicity around the start codon (Figure S2). Provided with small subunit profiling, the initiation rate (IR)— the number of 80S elongating ribosomes relative to 43S scanning^5^ — can also be calculated. Together with the sequence, this allows for investigating which initiation sequences lead to the most productive elongation (Figure 3D and 3E).

### Detecting translated upstream open reading frames

To illustrate an example of a search for novel translated regions we applied ORFik to the problem of discovering differential uORF usage across 3 developmental stages of zebrafish embryogenesis: 12 hours post-fertilization (hpf), 24 hpf, and 48 hpf^26^. Using public ribo-seq, RNA-seq^26^, and CAGE^48^ data we used ORFik to derive stage-specific 5’UTRs and searched these for all ATGs and near-cognate start codons. ORFik supports a number of features that have been shown to correlate with translation activity (Table S3). We used ORFik to calculate all these metrics per stage for 1) all uORF candidates, 2) all CDSs of known protein-coding genes and 3) random regions from the 3’ UTRs. We used these to train a random forest model on each stage with the H2O R-package^49^ using the CDSs as a positive set and the 3’UTR regions as the negative set (Supplementary Note 1). This model was used to predict translational activity over the uORF candidates resulting in 8,191 unique translated uORFs across all 3 stages (12hpf: 4,042, 24hpf: 2,178, 48hpf: 2,676). Of these, only 133 are translated in all stages (Figure 4A,B). Given the stringency of our training set which consists of long, translated proteins (positive) contrasted with short non-translated 3’UTR regions (negative) this should be considered a very conservative prediction. More nuanced predictions could be achieved by tuning these sets to better represent the typically shorter, more weakly translated uORFs.

**Figure 4:**
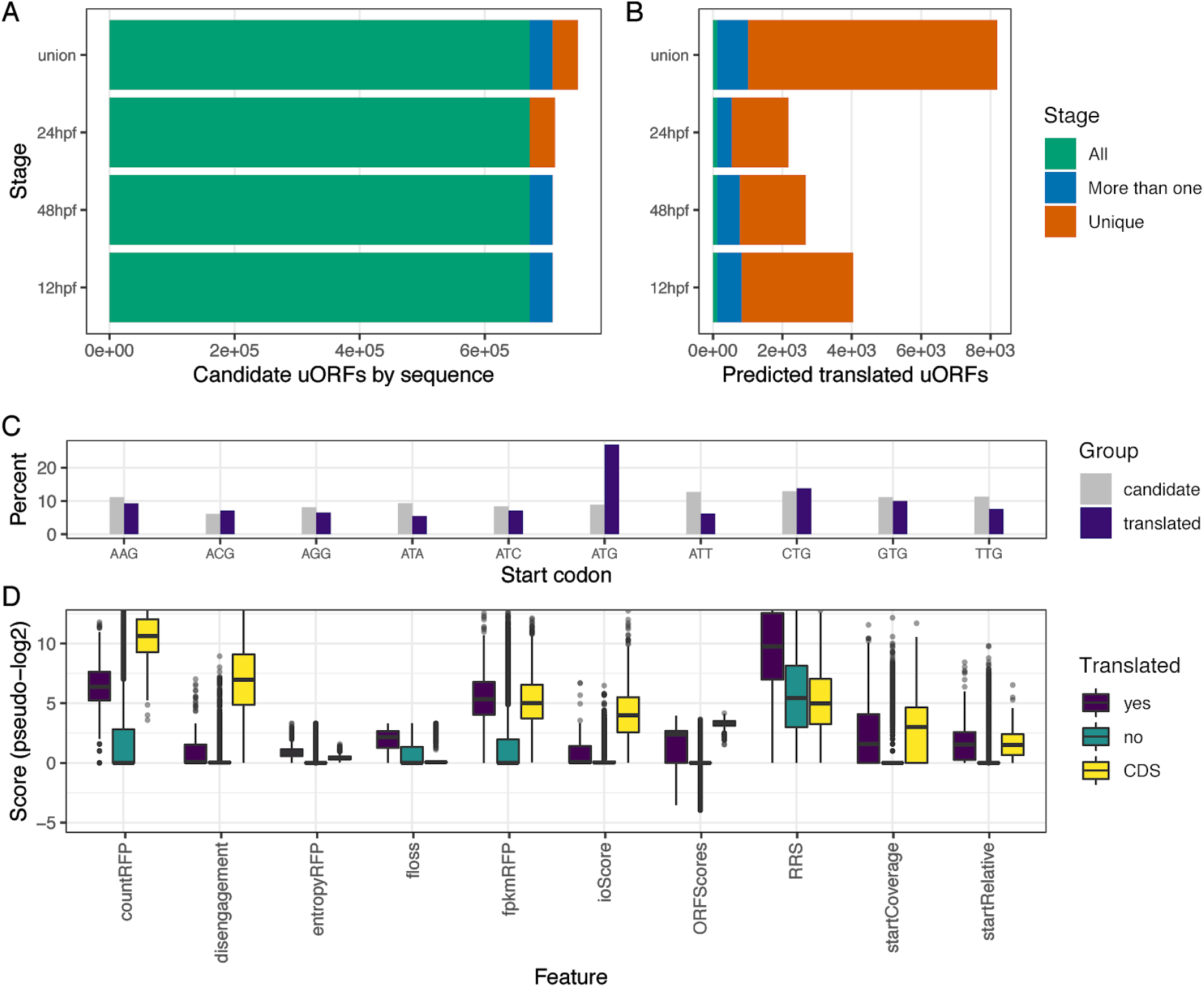
Prediction of uORFs in three zebrafish developmental stages. *A) Number of candidate uORFs identified from sequence per stage*. The *union group describes the union set of all Ihree timepoints. uORFs are colored whether they are common Io all stages (green), shared between multiple (blue), or unique to one stage (brown). B) Number of uORFs predicted t o be actively translated. Direct comparison of overlapping uORFs between stages can be seen in Figure S8. C) Distribution of start codons for candidate uORFs (gray) and predicted Iranslated uORFs (purple). D) Metrics used as Ieatures in classification (x-axis) shown for uORFs predicted Io be Iranslated (purple), predicted not to be translated (green), and protein-coding CDSs (yellow). See Table S3 for a description of features. Y-axis pseudo-log2 score (abs(log2(score)) if score > 0.01, −log2(-score) if score < −0.01, and 0 if absolute value of score is < 0.01)*.

For the predicted translated uORFs a clear shift in the usage of start codons can be seen relative to all possible candidate uORFs by sequence (Figure 4C). While purely sequence-predicted uORFs show a relatively uniform distribution in start codon usage, a clear bias towards ATG followed by CTG can be seen for the translated uORFs. Since the classifier favors uORFs that start with start codons known to be strong initiators, despite not using start codon sequence as a feature, suggest that it is able to identify uORFs that are actively translated. Since the classifier is trained on CDSs the resulting uORFs also have features that resemble known protein-coding genes (Figure 4D).

## Summary

In summary, we have developed ORFik, a new tool for streamlining analysis of ORFs and translation. ORFik introduces hundreds of tested, documented, and optimized methods to analyze and visualize ribosome coverage over transcripts. It supports a range of data formats and can be used to create complete pipelines from read processing and mapping to publication-ready figures. We demonstrate its use on a transcriptome-wide study of translation initiation and quick annotation of translated uORFs. Together, this empowers users to perform complex translation analysis with less time spent on coding, allowing the user to focus on biological questions.

## Availability and requirements

- **Project name:** *ORFik*
- **Project home page:** http://bioconductor.org/packages/ORFik
- **Archived version:** ORFik
- **Operating system(s):** Platform independent
- **Programming language:** R
- **Other requirements:** R version ≥3.6, Bioconductor version ≥3.10
- **License:** MIT + file LICENSE
- **Any restrictions to use by non-academics:** none

## Abbreviations

ORF: Open reading frame
CDS: Coding sequence (main ORF of mRNA)
5’ UTR: 5’ Untranslated region (also called transcript leader)
3’ UTR: 3’ Untranslated region (also called transcript trailer)
uORF: Upstream open reading frame
NGS: Next-generation sequencing
RFP: Ribosome-protected footprint (also referred to as RPF)
P-shifting: Relative 5’ end shifting of Ribo-seq reads to their estimated P-site
HPF: Hours post-fertilization
TSS: Transcription start site
TIS: Translation initiation site
IGV: Integrative genomics viewer
SSU: Small subunit (ribosomal)
LSU: Large subunit (ribosomal)
80S: 80S elongating Ribosome
40S/43S: 40S/43S Ribosomal preinitiation complex

## Acknowledgements

We would like to thank the Bioconductor core team and community for providing valuable feedback for developing *ORFik*.

## Authors’ contributions

KL, HT and EV conceived the idea for ORFik. HT, MS and KL wrote the code. YTC, KC and EV have contributed to the code and functionality. All authors wrote and approved the final manuscript.

## Consent for publication

Not applicable.

## Ethics approval and consent to participate

Not applicable.

## Supplementary

Code for figures, tables and alignment of data: https://github.com/Roleren/ORFik_article_code

### Supplementary Tables

**Table S1:**
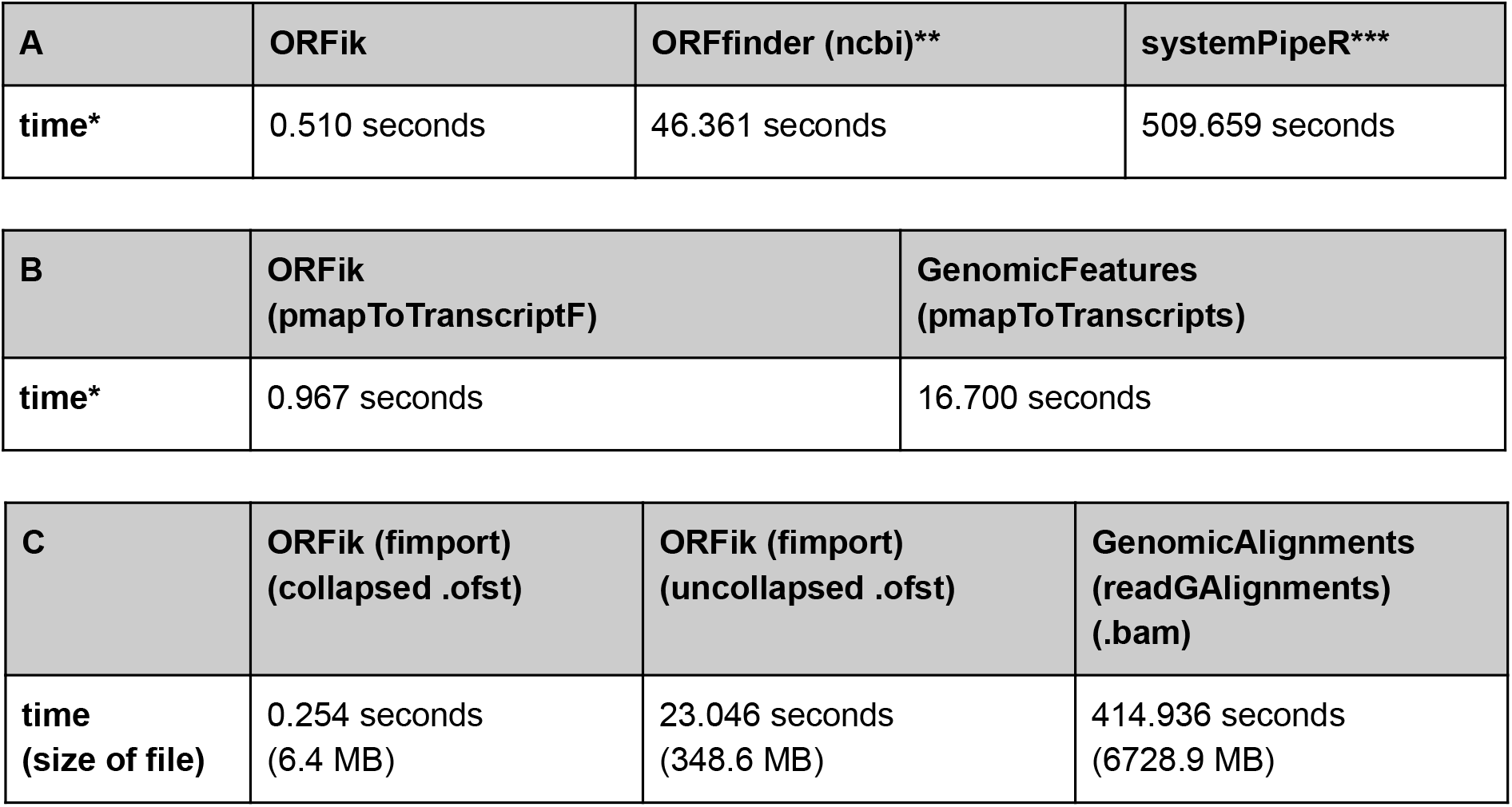
Runtime comparison of ORFik versus other tools. ***A) Comparison of runtime for tools finding ORFs*.** *Finding uORFs on all 5’ UTRs in zebrafish Iranscriptome (Ensembl Danio rerio GRCz10, using a total of 11,343 5’ UTRs)*. *Arguments given: minimum length ORF: 30 bases (ORFfinder’s minimum), strands: both, start codons: ATG, stop codons: TAA, TAG, TGA. Longest ORF per stop codon: FALSE*. *\*Average time of 3 runs. **ORFfinder linux x64 desktop version 2020-03-11 13:30^50^. ***It should be noted that* systemPipeR::predORF was not designed for large scale ORF detection. Version: 1.23.4 ***B) Comparison in runtime of genomic to transcript coordinate mapping*** *Mapping all transcripts Io Iranscript coordinates (Ensembl Danio rerio GRCz10, using a Iotal of 57,369 transcripts). Output of Iunctions are here identical. *Average time of 3 runs. GenomicFeatures version: 1.42.1* ***C) Comparison in load time of.bam file vs .ofst using ORFik::fimport.** Three different file types that load identical information into R (except QNAMES which are lost when collapsing reads), showing Ihe increase in speed for loading collapsed. ofst files as compared Io bam files. File used: 2hpf sample of ribo-seq from Bazzini et al 2014, see Table S5*.

| A | ORFik | ORFfinder (ncbi)** | systemPipeR*** |
| --- | --- | --- | --- |
| time* | 0.510 seconds | 46.361 seconds | 509.659 seconds |

| B | ORFik<br>(pmapToTranscriptF) | GenomicFeatures<br>(pmapToTranscripts) |
| --- | --- | --- |
| time* | 0.967 seconds | 16.700 seconds |

| C | ORFik (fimport)<br>(collapsed .ofst) | ORFik (fimport)<br>(uncollapsed .ofst) | GenomicAlignments<br>(readGAlignments)<br>(.bam) |
| --- | --- | --- | --- |
| time<br>(size of file) | 0.254 seconds<br>(6.4 MB) | 23.046 seconds<br>(348.6 MB) | 414.936 seconds<br>(6728.9 MB) |

**Table S2:** Coverage transformations in ORFik. The coverage transformation available in ORFik through the function coverageScorings. This tunction can calculate meta coverage (summarizing over multiple regions). It takes as input the raw coverage per nucleotide per gene per fraction (e.g. mRNA-seq, Ribo-seq, read length) per feature (e.g. 5’ UTRs, CDS, 3’ UTRs) as input. The output is either a normalization of the data, a grouping or a combination of these two.

| Coverage transformations | Description: |
| --- | --- |
| zscore | $(x - \text{mean counts}) / \text{standard deviation of counts}$ |
| transcriptNormalized | $x / \text{total counts of region (example: fraction of reads mapping to a transcript)}$ |
| mean | mean x |
| median | median x |
| sum | sum x |
| log2sum | $\log_2 (\text{sum } x)$ |
| log10sum | $\log_{10} (\text{sum } x)$ |
| sumLength | $x / \text{number of regions}$ |
| meanPos | mean x / binned window size |
| sumPos | sum x / binned window size |
| frameSum | sum x (per frame) |
| frameSumPerL | sum x (per frame per read length) |
| frameSumPerLG | sum x (per frame per read length per gene) |
| fracPos | fraction of counts per nucleotide (separate by gene) |
| periodic | Does x show a periodicity of 3: (TRUE / FALSE) |
| NULL | Input returned directly |

**Table S3:**
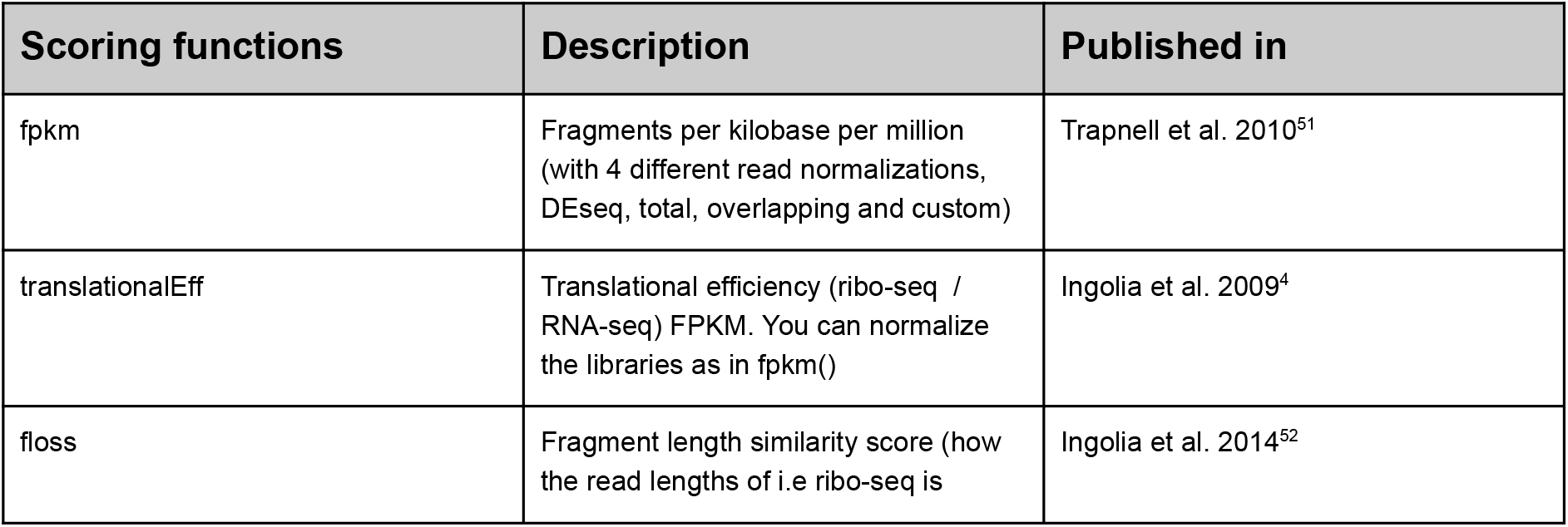
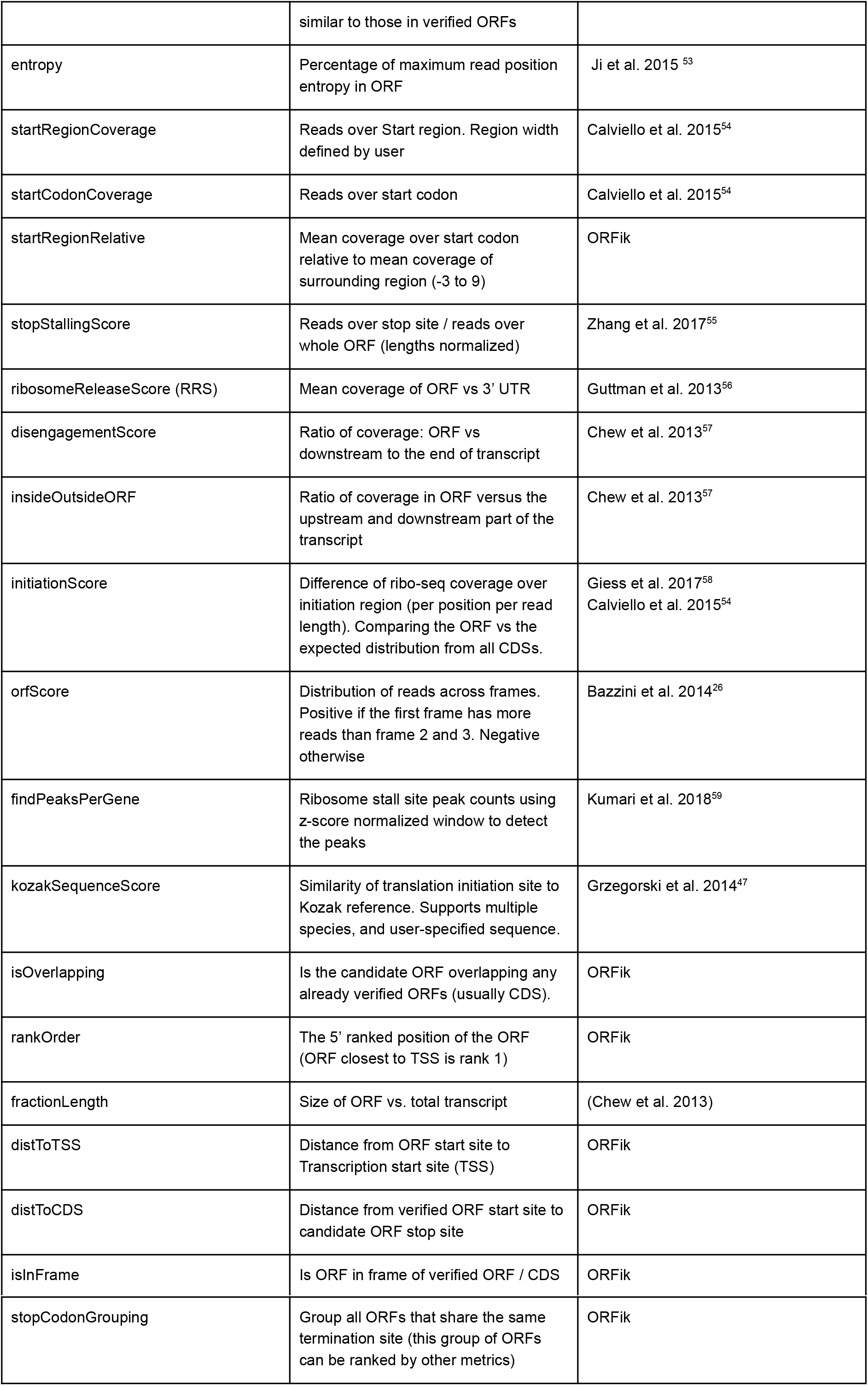
Scoring functions included in ORFik. ORFik supports many published scoring functions for prediction of translated ORFs. The scoring functions column shows the names of the sequence and translation features used in the computeFeatures function and a few stand alone functions. Description column briefly describes each feature. Using ORFik back-end functions, you can easily create your own functions if needed.

**Table S4:** H2O random forest prediction model statistics. ***A:** Consistency over repeated runs*. To show that the pipeline’s non-deterministic training step does not significantly impact the result, the uORFs from zebrafish development-stages were run through the pipeline 3 times, shown here in true positive rate and true negative rate. Giving the number of matched predicted true (translated) and negative (not translated) between the 3 runs. **B:** 10-fold cross validation to determine training model statistics. Shown are average accuracy, precision and recall of random forest training models over all 3 zebrafish development-stages.

| A: Scoring feature | Description: |
| --- | --- |
| true positive rate (match in predicted positives) | 98.1% |
| true negative rate (match in predicted negatives) | 99.5 % |
| B: Scoring feature | Description: |
| accuracy (mean +/- sd) | 0.9974 +/- 0.0012 |
| precision (mean +/- sd) | 0.9753 +/- 0.0102 |
| recall (mean +/- sd) | 0.9878 +/- 0.0073 |
| prediction type | Binary class prediction |
| number of trees (training) | 100 |

**Table S5:** Data used in this manuscript. RNA-seq, ribo-seq and RCP-seq data was trimmed with fastp and aligned with STAR using ORFik wrapper. See the ORFik STAR and fastp script for the default arguments not specified here. For full details of data used see the attached alignment scripts.

| Feature | Ribo-seq | RNA-seq | CAGE | TCP-seq | CAGE |
| --- | --- | --- | --- | --- | --- |
| Reference | Bazzini et al 2014 (GSE53693) | Bazzini et al 2014 (GSE53693) | Nepal et al 2013 (SRA055273) | Bohlen et al 2020 (GSE139132) | Forrest et al, FANTOM5 Consortium 2014 (DRR041459) |
| Species | Danio rerio | Danio rerio | Danio rerio | Homo sapiens | Homo sapiens |
| Genome-annotation | Danio_rerio.GRCz10 (ensembl) | Danio_rerio.GRCz10 (ensembl) | Danio_rerio.GRCz10 (ensembl) | GRCh38.101 (ensembl) | GRCh38.101 (ensembl) |
| Alignment, run mode: | single end | single end | single end | single end | single end |
| **Adapter**<br>**sequence**<br>**trimmed (3')** | AGATCGGAAGA<br>GC | AGATCGGAAGAGC | - | AGATCGGAA<br>GAGC | TGGAATTCT<br>CGG |
| **3' end bases**<br>**trimmed** | 0 | 3 | 1 | 0 | 0 |
| **minimum**<br>**length** | 20 | 20 | 20 | 20 | 20 |

**Table S6:** Comparison of functionality from translation tools. Figure updated and expanded from Ribotoolkit manuscript supplementary data (Table S3)^11^.

| Feature | ORFik | RiboToolkit | Ribo-seQC | RiboTools | riboSeqR | plastid | RiboProfiling | RiboGalaxy | Proteoformer | systemPipeR | RiboVIEW | riboStreamR | RiboFlow/<br>RiboR/RiboPy |
| --- | --- | --- | --- | --- | --- | --- | --- | --- | --- | --- | --- | --- | --- |
| NGS Input files |  |  |  |  |  |  |  |  |  |  |  |  |  |
| FASTA | ✓ | ✓ |  |  |  |  |  |  |  |  |  |  |  |
| FASTQ | ✓ |  |  |  |  |  |  | ✓ | ✓ | ✓ |  |  | ✓ |
| BAM | ✓ |  | ✓ | ✓ | ✓ | ✓ | ✓ | ✓ |  | ✓ | ✓ | ✓ |  |
| Link (URL) |  | ✓ |  |  |  |  |  |  |  |  |  |  |  |
| Project ID (SRA) | ✓ |  |  |  |  |  |  |  |  |  |  |  |  |
| NGS input types (* can also align type) |  |  |  |  |  |  |  |  |  |  |  |  |  |
| Ribo-seq | ✓* | ✓* | ✓ | ✓ | ✓ | ✓ | ✓ | ✓* | ✓* | ✓* | ✓ | ✓ | ✓* |
| mRNA-seq | ✓* | ✓ | ✓ |  | ✓ | ✓ |  | ✓* | ✓* | ✓* |  |  | ✓* |
| TCP / RCP-seq | ✓* |  | ✓ |  |  |  |  |  |  |  |  |  |  |
| CAGE | ✓* |  |  |  |  |  |  |  |  |  |  |  |  |
| Quality control |  |  |  |  |  |  |  |  |  |  |  |  |  |
| Contamination QC | ✓ | ✓ | ✓ |  |  | ✓ |  |  | ✓ | ✓ | ✓ | ✓ | ✓ |
| Alignment QC | ✓ |  |  |  |  |  |  | ✓ | ✓ | ✓ |  |  | ✓ |
| Annotation QC | ✓ | ✓ | ✓ | ✓ | ✓ | ✓ | ✓ | ✓ | ✓ | ✓ | ✓ | ✓ | ✓ |
| Analysis QC | ✓ | ✓ | ✓ | ✓ | ✓ | ✓ | ✓ | ✓ | ✓ | ✓ | ✓ | ✓ | ✓ |
| Active ORF translation |  |  |  |  |  |  |  |  |  |  |  |  |  |
| Active CDS calling | ✓ | ✓ |  |  | ✓ |  |  |  | ✓ | ✓ |  |  |  |
| Active uORF calling | ✓ | ✓ |  |  |  |  |  |  | ✓ | ✓ |  |  |  |
| Active dORF calling | ✓ | ✓ |  |  |  |  |  |  |  |  |  |  |  |
| Active prokaryote ORF calling (circular genomes) | ✓ |  |  |  |  |  |  |  |  |  |  |  |  |
| Expression and translational analyses |  |  |  |  |  |  |  |  |  |  |  |  |  |
| mRNA expression | ✓ | ✓ | ✓ |  | ✓ | ✓ |  | ✓ |  | ✓ |  | ✓ |  |
| RFP (80S) expression | ✓ | ✓ | ✓ |  | ✓ | ✓ | ✓ | ✓ | ✓ | ✓ |  | ✓ |  |
| 40S expression | ✓ |  | ✓ |  |  |  |  |  |  |  |  |  |  |
| TE analysis | ✓ | ✓ |  |  | ✓ | ✓ |  | ✓ |  | ✓ |  |  |  |
| IR analysis | ✓ |  |  |  |  |  |  |  |  |  |  |  |  |
| Differential analysis | ✓ | ✓ |  |  | ✓ |  |  | ✓ |  | ✓ |  |  |  |
| Other functionalities |  |  |  |  |  |  |  |  |  |  |  |  |  |
| Metagene plots | ✓ | ✓ | ✓ | ✓ | ✓ | ✓ | ✓ | ✓ |  | ✓ | ✓ | ✓ | ✓ |
| Heatmap plots | ✓ |  | ✓ |  |  |  |  |  |  |  |  |  |  |
| Codon analysis plots | ✓ | ✓ | ✓ |  | ✓ | ✓ | ✓ | ✓ | ✓ | ✓ | ✓ |  |  |
| P-site analysis | ✓ | ✓ | ✓ | ✓ | ✓ | ✓ | ✓ | ✓ | ✓ | ✓ | ✓ | ✓ | ✓ |
| Customizable |  |  |  |  |  |  |  |  |  |  |  |  |  |
| Customizable | ✓ | ✓ | ✓ | ✓ | ✓ | ✓ | ✓ | ✓ | ✓ | ✓ | ✓ | ✓ | ✓ |
| Fully customizable backend | ✓ |  |  |  |  |  |  |  |  | ✓ |  |  |  |

### Supplementary Figures

**Figure S1:**
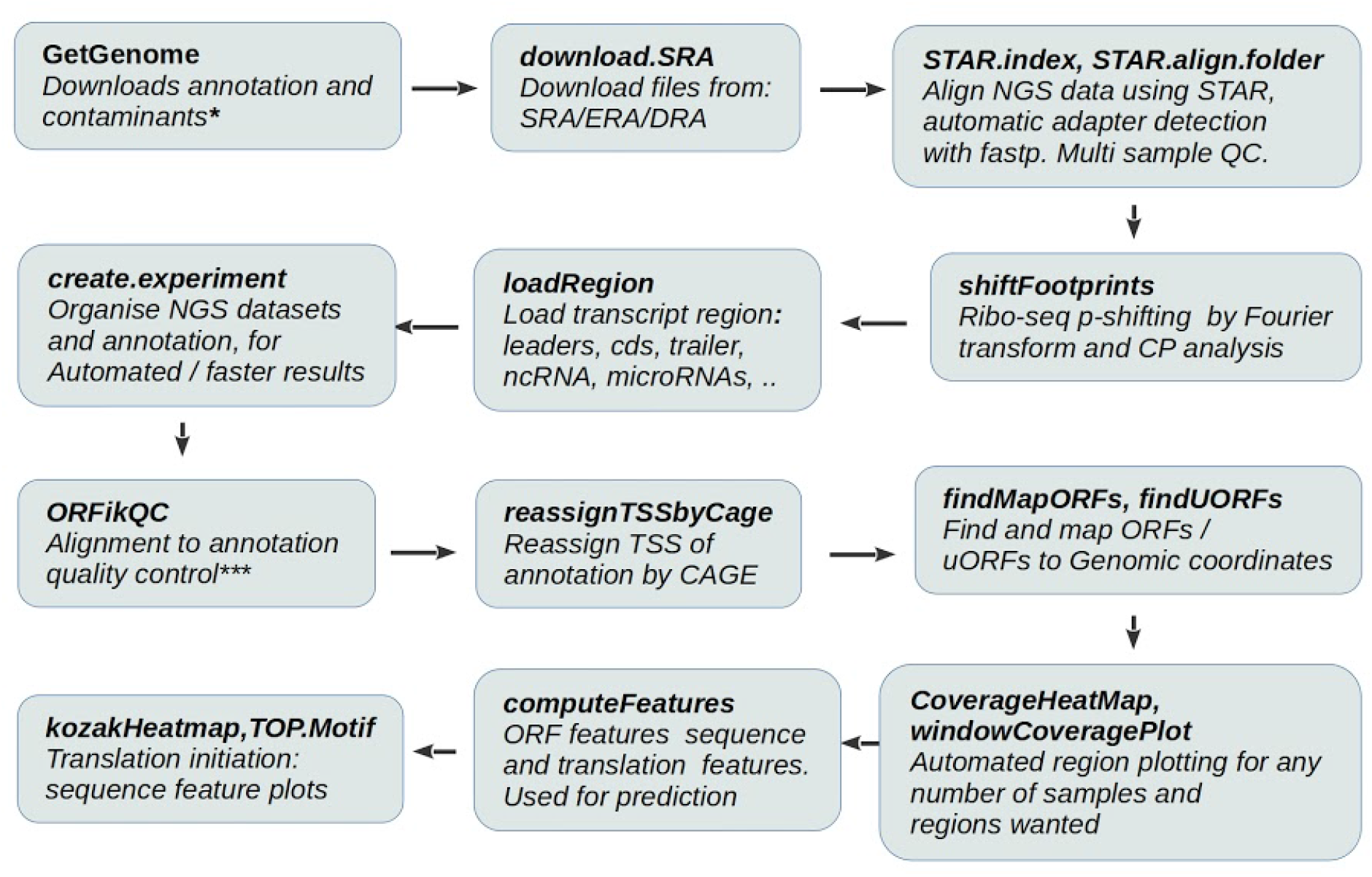
Example of ORFik workflow. ORFik simplifies and automates all data handling steps, from downloading genome, annotations and experimental data through p-shifting and mapping up to novel ORF detection and classification. * Can download contaminant sequences for depletion in NGS libraries, tike Illumina Phix genome or noncoding RNAs. **Gives direct access to NGS library locations, QC report etc. *** QC report optimized (but not exclusive) for STAR aligner^32^ and DESeq2^41^.

**Figure S2:**
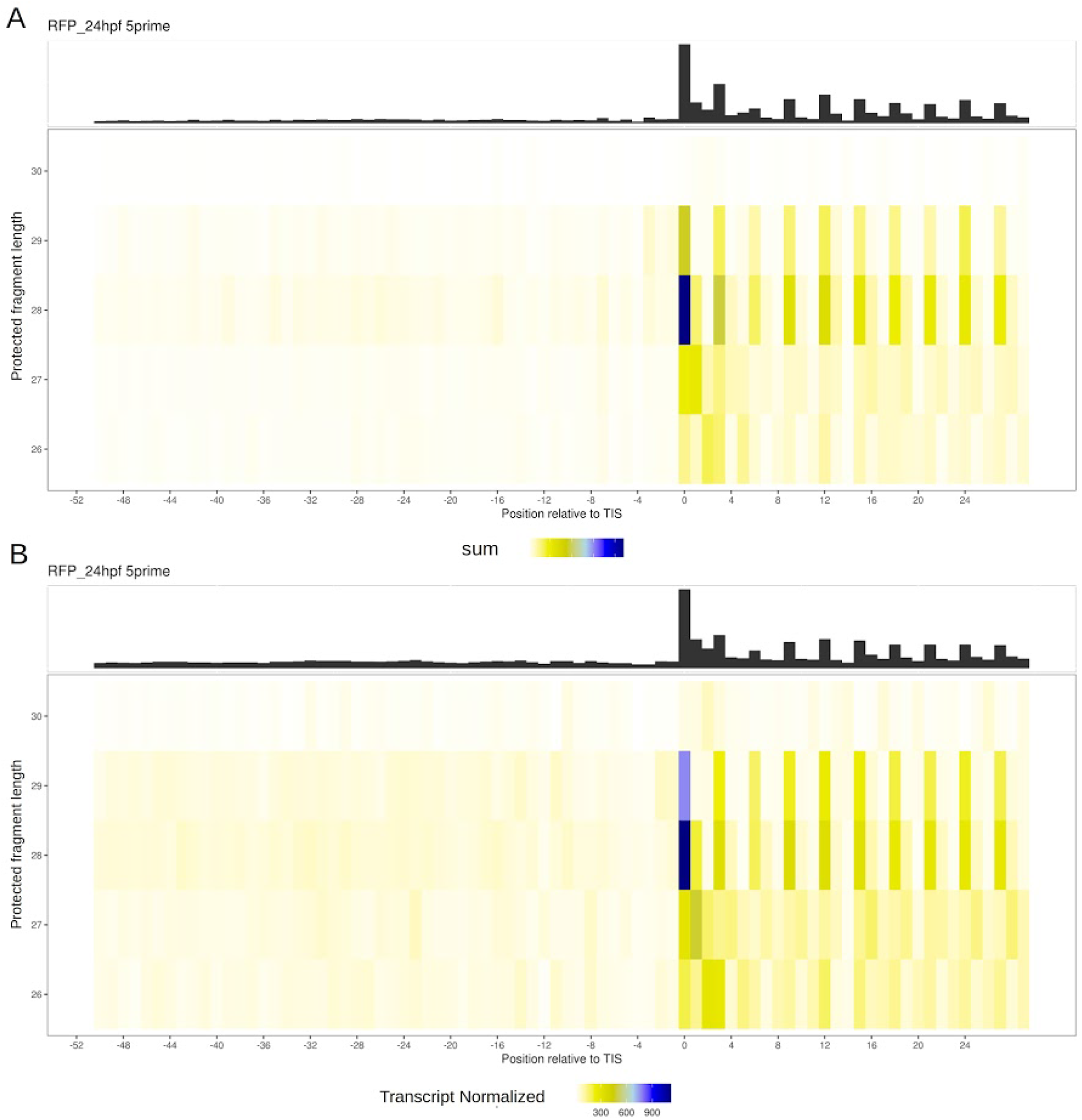
Meta coverage heatmap around TIS of P-site shifted ribo-seq data using different read count normalization. y-axis: Read lengths 26 to 30. x-axis: nucleotide position around TIS (−52 to +29). Colors show A) count of 5’ ends of reads and B) transcript-normalized counts (all counts for one window sum to 1). Data from Bazzini et al 2014 ribo-seq (Table S5).

**Figure S3:**
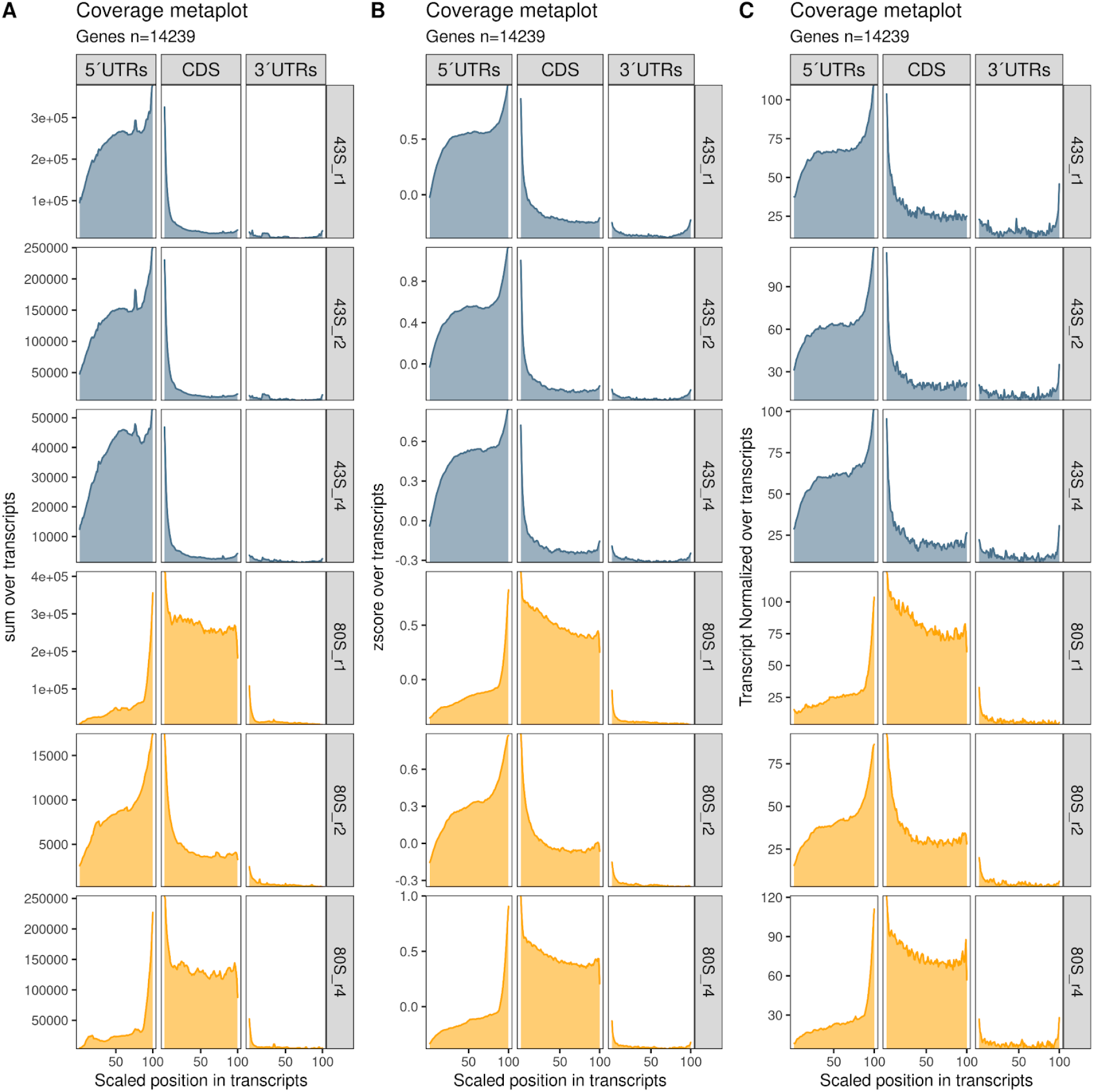
Meta coverage over mRNAs in non-normalized and normalized counts split by region. Transcripts are split into 5’ UTRs, CDSs and 3’ UTRs. The x-axis shows the relative position normalized to a length of 100 and y-axis shows score value. The scores are **A:** Sum (raw counts), **B:** z-score (the mean of the z-scores from all transcripts at that position), **C:** transcript normalized (counts per transcript sum to 1). Colors describe library type; orange (80S) and blue (43S). Data from TCP-seq in Bohlen et al 2020 (Table S5).

**Figure S4:**
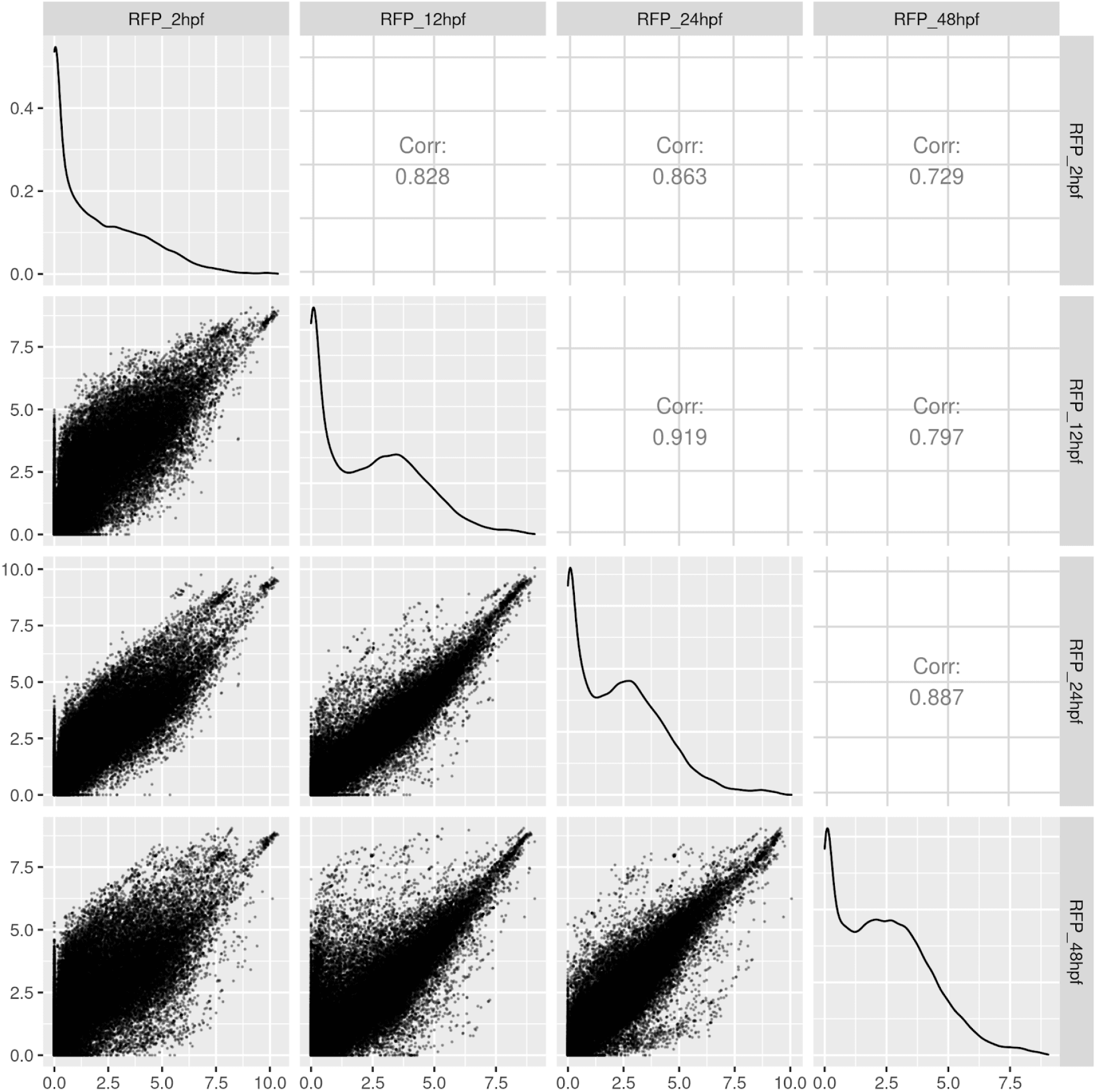
Correlation plots between all samples. Log2 FPKM correlation of genes between the 4 stages used of ribo-seq from Bazzini et al 2014. This is part of the default QC.

**Figure S5:**
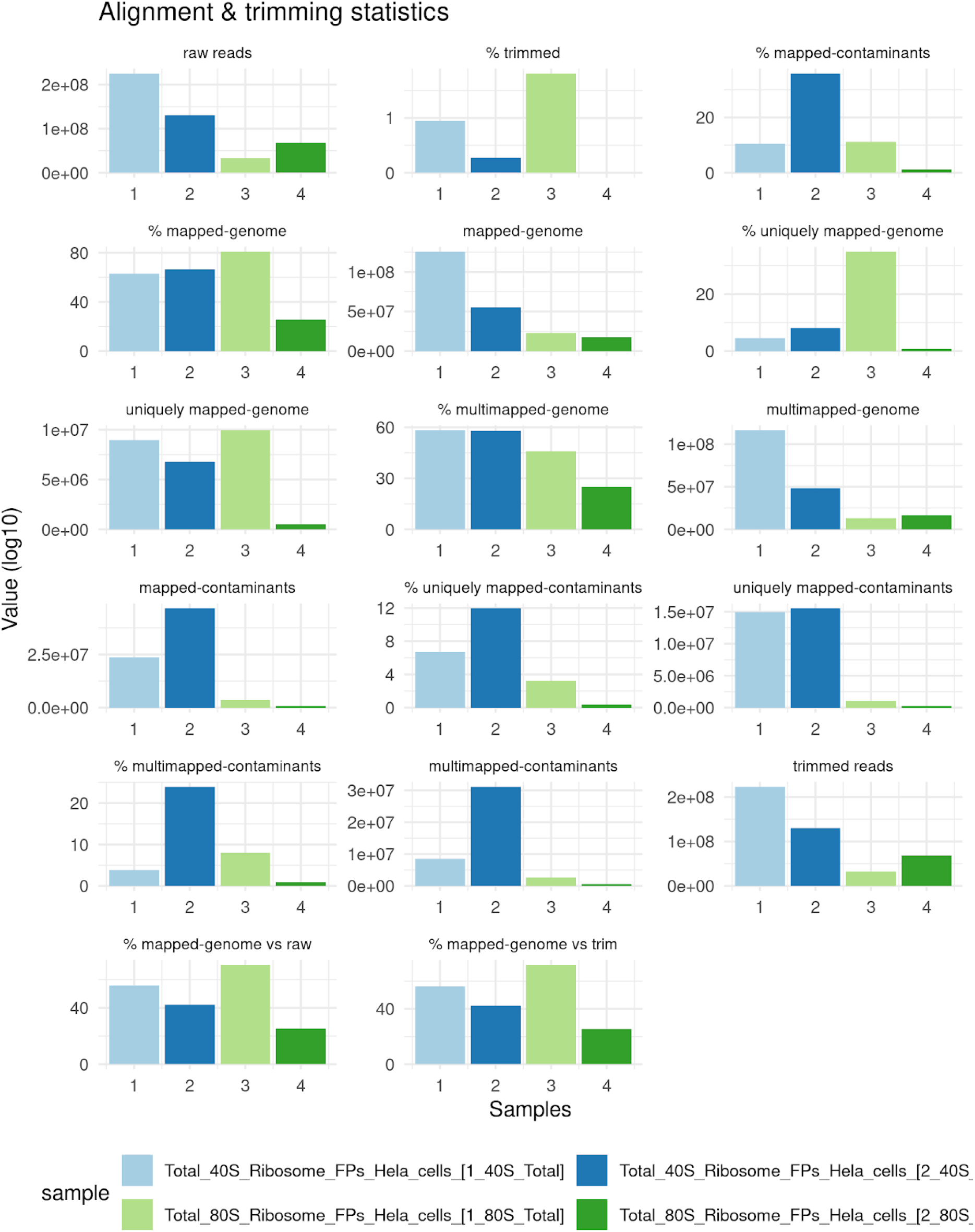
Alignment and trimming statistics. Extracted and visualized output from fastp preprocessing and STAR alignments. Shown are the amount of reads in counts and percentage (%) at the following steps: raw reads from sequencing, trimmed reads (from fastp), contamination depletion (here defined as phix, rRNA, iRNA and non coding RNAs), genome alignment (from STAR). Users will in addition get more detailed FASTQ read statistics from fastp, among others: quality scores, adapter analysis, duplication rates, base content ratios and k-mer counting. Example outputs from fastp can be found here: http://opengene.org/fastp/fastp.html Data from 4 samples of TCP-seq from Bohlen et al 2020.

**Figure S6:**
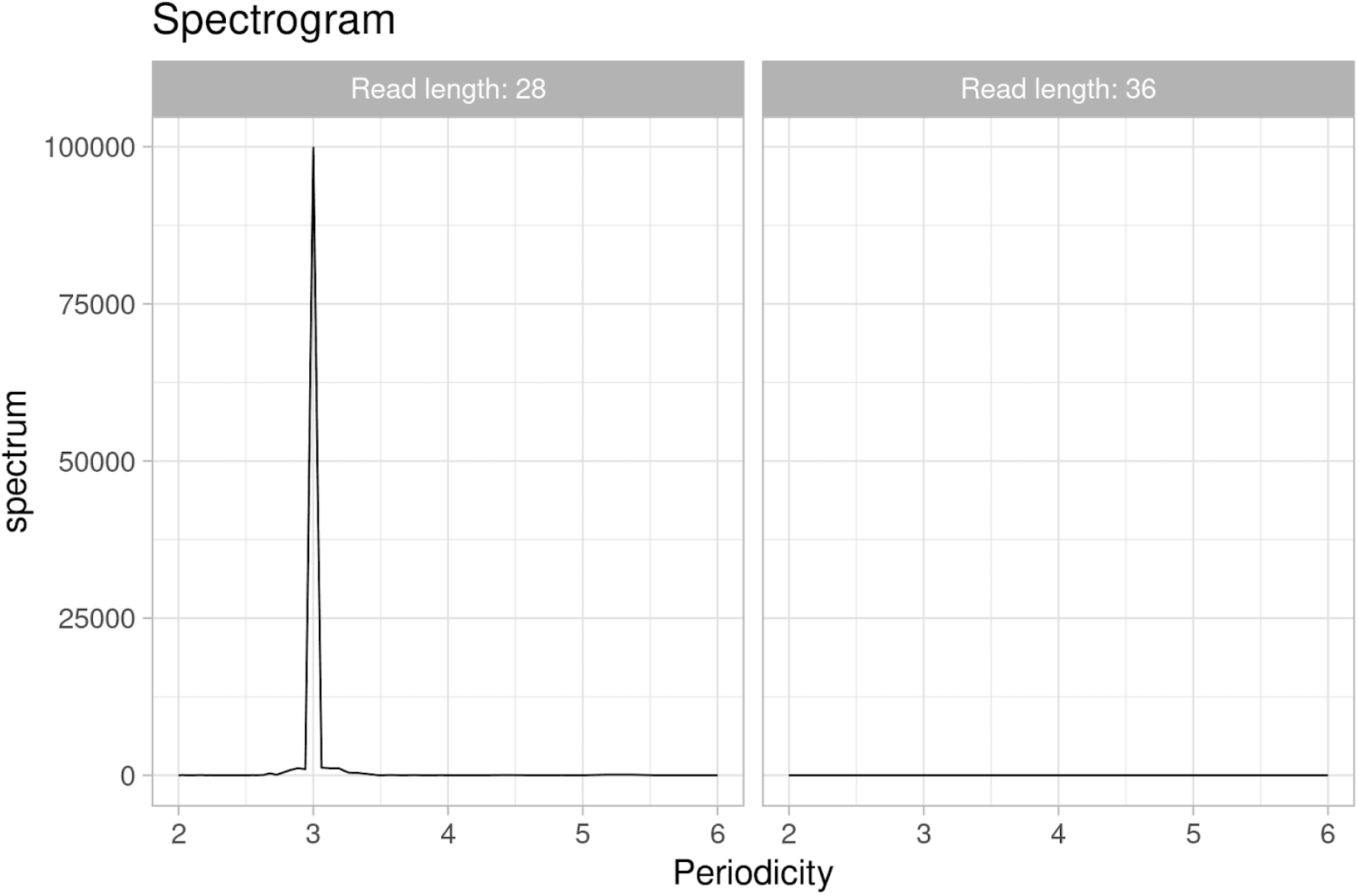
Spectrogram analysis of Ribo-seq read lengths. Spectrograms showing periodicity for two read lengths. The shorter (28nt) shows a 3nt-periodicity, while the longer (36nt) does not. Read lengths that do not have a 3 nt read length periodicity will be filtered out for 80S libraries. The peak must be based on at least 1000 reads across the search regions. Data from 2 hpf from Bazzini et al 2014 (Table S3).

**Figure S7:**
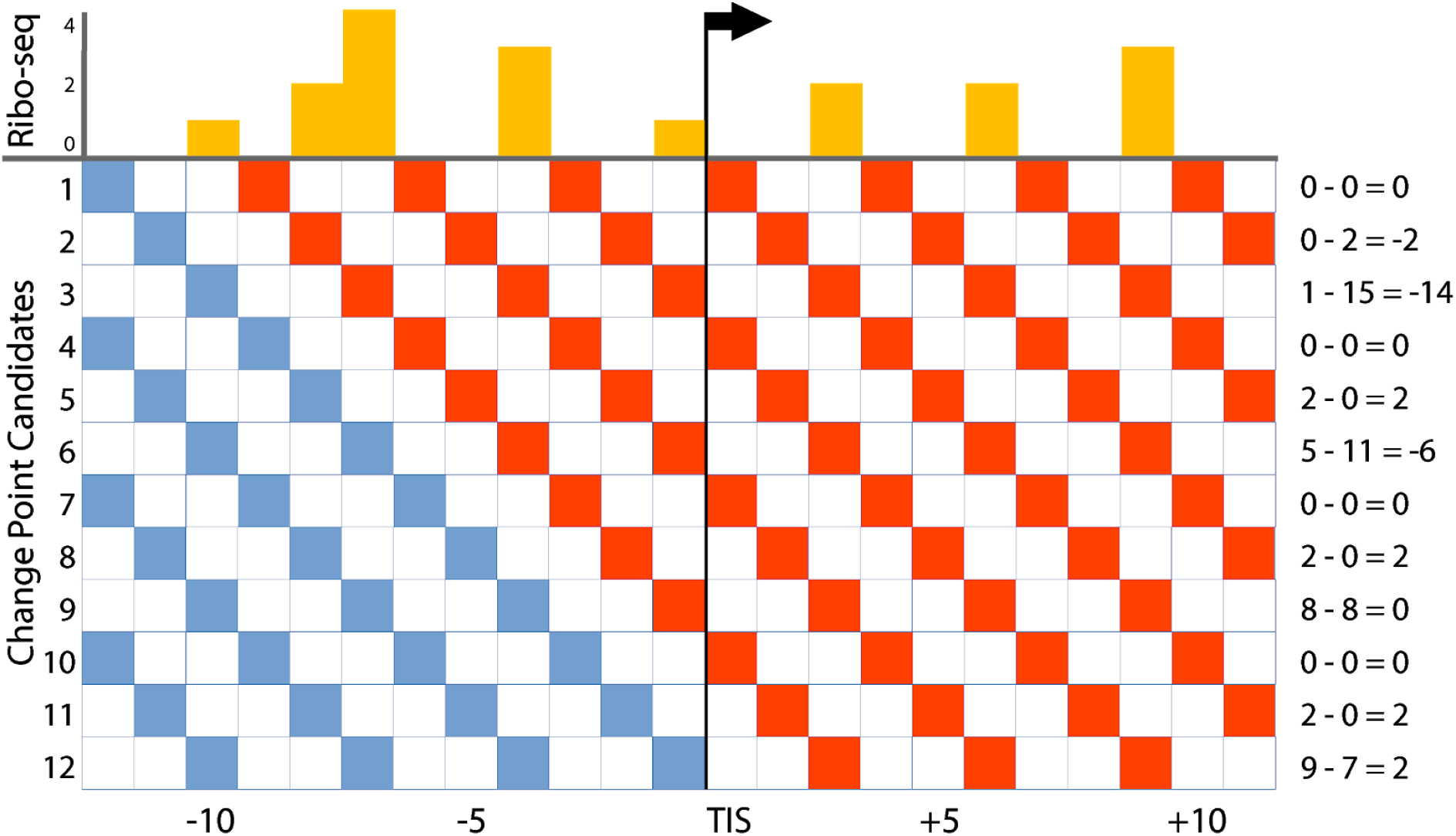
Graphical explanation of change point analysis during P-site offset-detection in ORFik. Separately for every read length, the 5’ ends of all reads are summed up tn a window (+/- 30nt) around the translation initiation site (TIS). From this distribution (yellow) every position tn the window (x-axis) is evaluated as a potential change point (y-axis). For each position the sum of the reads tn the same translational frame (every 3rd nucleotide) tn a window upstream (blue squares) of this is compared Io the sum of reads at the position and downstream (red squares). These sums are given tn the right column. The change point with the largest difference, tn this case #3 with −14, is selected and the shift is calculated based on the distance between this point and the TIS, tn this case 7 nts.

**Figure S8:**
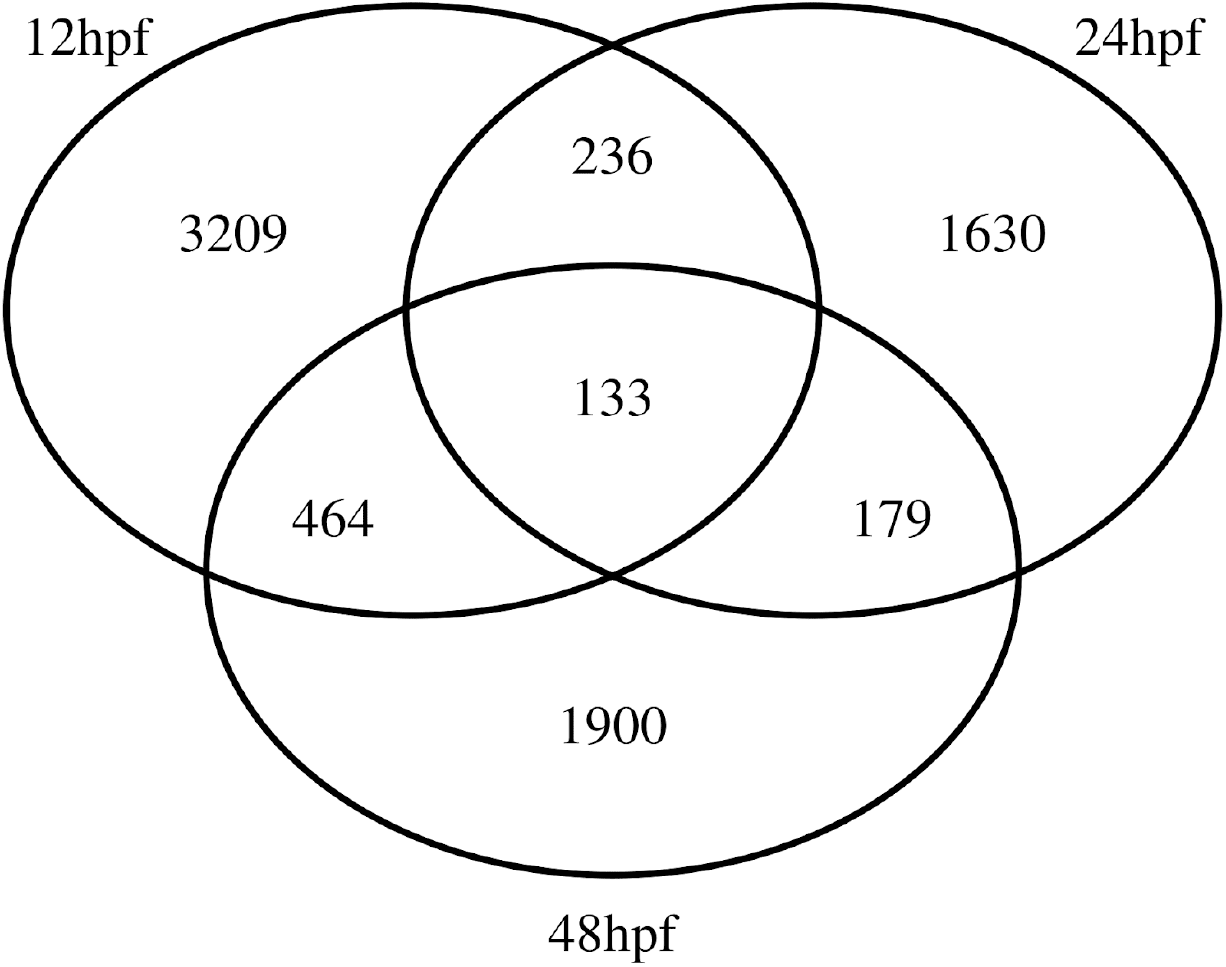
Predicted uORFs across stages. Venn diagram showing uORFs predicted between stages tn the uORF prediction.

**Supplementary Note 1**: Detailed description of uORF prediction pipeline

1. Determine the search-space for uORFs for each CAGE library. The search-space is defined as CAGE derived 5’UTRs + CDS. If CAGE libraries are not included, all CAGE steps are skipped.
2. Find all uORFs in the search-space with default options: Start codons: “ATG|CTG|TTG|GTG|AAG|AGG|ACG|ATC|ATA|ATT” Stop codons: “TGA|TAA|TAG” Minimum length: 6 bases (start and stop codon) Longest ORF per stop codon: FALSE (all uORFs per stop codon included). Requirements: Candidate uORF must start in the 5’ UTR and end the base before the CDS stop site. Candidate uORF can’t be in frame relative to the CDS, if the uORF overlaps the CDS.
3. Create unique identifiers for each candidate uORF found. This identifier is defined as transcript id + chromosome + candidate uORF start location + width, separated with an underscore.
4. Find the sequence and NGS features of candidate uORFs using ORFik::computeFeatures function.
5. Store all uORF features in a database (RMysql) for persistence.
6. Create 1 training model and prediction model per stage / cell line. Replicates of the same stage will be grouped together, mean score value over replicates.
7. To create the positive training data, the most highly translated CDS are selected (see below), while for the negative set random untranslated windows in the 3’UTRs are sampled (see below).

a. For the positive set (CDS) this filtering is applied: (Ribo-seq FPKM > 1 & FPKM > 25th percentile) and (counts > 10 and counts > 25 percentile) and (start codon coverage > 75th percentile) and (periodicity ORFscore > 1). This will train the model conservatively on CDSs that have high coverage around the start codon and a clear periodicity in the Ribo-seq.
b. For the negative set (3’ UTRs) this filtering is applied: (Ribo-seq FPKM < 75th percentile) or (coverage < 75 percentile) or (start codon coverage < 75th percentile) or (periodicity ORFscore < 0.5). This presents the model with a diverse data set of non-translated sequences that either have little or no coverage, or coverage from overlapping translated ORFs.
8. Assign CDSs that did not pass filters into the negative set.
9. Train the random forest model on the positive and negative sets (Table S4).
10. Predict uORFs using the model on candidate uORFs with their respective feature data.

Run time for the uORFome pipeline for the data shown in Figure 4 was 14 minutes +/- 2 minutes over 3 runs on CentOS 7 having 196 cores intel with 2 TB memory. Pipeline used between 1-48 cores depending on the different parts of the pipeline.

